# Shedding light on the control of CatSper Ca^2+^ channels by cAMP and chemicals used to probe cAMP signaling

**DOI:** 10.1101/2020.02.10.942029

**Authors:** Tao Wang, Samuel Young, Frank Tüttelmann, Albrecht Röpke, Claudia Krallmann, Sabine Kliesch, Xu-Hui Zeng, Christoph Brenker, Timo Strünker

## Abstract

The sperm-specific CatSper channel controls the influx of Ca^2+^ into the flagellum and, thereby, the swimming behavior of sperm. A hallmark of human CatSper is its polymodal activation by membrane voltage, intracellular pH, and oviductal hormones. Whether CatSper is also activated by signaling pathways involving an increase of cAMP and ensuing activation of protein kinase A (PKA) is, however, a matter of controversy. Here, using kinetic ion-sensitive fluorimetry and patch-clamp recordings, we study transmembrane Ca^2+^ flux and membrane currents in human sperm from healthy donors and from patients that lack functional CatSper channels. We show that human CatSper is neither activated by intracellular cAMP directly nor indirectly by the cAMP/PKA-signaling pathway. Moreover, we demonstrate that non-physiological concentrations of cAMP and membrane-permeable cAMP analogs used to mimic the action of intracellular cAMP activate human CatSper from the outside via a previously unknown extracellular cyclic nucleotide-binding site. Finally, we demonstrate that the effects of common PKA inhibitors on human CatSper rest on off-target drug actions on CatSper itself rather than on inhibition of PKA. We conclude that the concept of an intracellular cAMP/PKA-activation of CatSper is primarily based on unspecific effects of chemical probes used to interfere with cAMP signaling. Altogether, our findings solve several controversial issues, which has important bearings on future studies of cAMP and Ca^2+^ signaling and the ligand-control of CatSper in sperm.

## Introduction

The CatSper channel (cation channel of sperm) represents the principal pathway for Ca^2+^ entry into the flagellum of sperm from many species (Kirichok et al., 2006; Lishko et al., 2011; Loux et al., 2013; Ren et al., 2001; Seifert et al., 2015; Strunker et al., 2011; Sumigama et al., 2015). The activity of CatSper is controlled by both the membrane potential (V_m_) and intracellular pH (pH_i_) (Hwang et al., 2019; Kirichok et al., 2006; Lishko et al., 2010; Lishko et al., 2011; Seifert et al., 2015; Strunker et al., 2011), and, in human sperm, also by oviductal steroids and prostaglandins (Brenker et al., 2018b; Lishko et al., 2011; Luo et al., 2019; Schiffer et al., 2020; Smith et al., 2013; Strunker et al., 2011; Williams et al., 2015). Thereby, CatSper translates changes of the chemical microenvironment into changes of [Ca^2+^]_i_ and swimming behavior, which enables sperm to reach the site of fertilization, to overcome the egg’s protective vestments, and, ultimately, to fertilize the egg (Achikanu et al., 2018; Alasmari et al., 2013; Oren-Benaroya et al., 2008; Ren et al., 2001; Rennhack et al., 2018; Schiffer et al., 2020; Tamburrino et al., 2014; Tamburrino et al., 2015). CatSper is, hence, absolutely required for fertilization in mice and humans (Avenarius et al., 2009; Avidan et al., 2003; Luo et al., 2019; Qi et al., 2007; Ren et al., 2001; Schiffer et al., 2020; Williams et al., 2015; Zhang et al., 2007).

Not only the control of [Ca^2+^]_i_ by CatSper, but also the flagellar cAMP dynamics is key for the function of sperm (Akbari et al., 2019; Balbach et al., 2018; Buffone et al., 2014; Esposito et al., 2004; Visconti et al., 1995). In mammalian sperm, cAMP is predominately synthesized by the soluble adenylyl cyclase (sAC) that is controlled by bicarbonate (Brenker et al., 2012; Buffone et al., 2014; Hess et al., 2005; Kleinboelting et al., 2014; Wennemuth et al., 2003b; Xie et al., 2006). Bicarbonate-induced synthesis of cAMP by sAC activates protein kinase A (PKA) (Buffone et al., 2014; Moseley et al., 2005); and the cAMP/PKA-signaling pathway controls the flagellar beat frequency and capacitation (Esposito et al., 2004; Hess et al., 2005; Morgan et al., 2008; Xie et al., 2006), a maturation process that primes sperm to fertilize the egg (Yanagimachi, 1994).

The interplay of Ca^2+^ and cAMP in sperm is only ill-defined. In particular, it has remained controversial whether intracellular cAMP and/or activation of PKA stimulate Ca^2+^ influx via CatSper. It is unequivocal that membrane-permeable analogues of cAMP (e.g. 8-Br-cAMP) that are used to mimic the action of intracellular cAMP activate CatSper in both mouse and human sperm (Brenker et al., 2012; Orta et al., 2018; Ren et al., 2001; Xia et al., 2007). However, in a series of studies by independent groups, elevation of intracellular cAMP levels in mouse and human sperm by bicarbonate or other measures, including adenosine, synthetic adenosine- or catecholamine-receptor agonists, photorelease of cAMP from caged cAMP, or control of cAMP by optogenetics, failed to stimulate Ca^2+^ influx via CatSper (Brenker et al., 2012; Carlson et al., 2007; Carlson et al., 2003; Jansen et al., 2015; Nolan et al., 2004; Schuh et al., 2006; Strunker et al., 2011; Wennemuth et al., 2003a). These results indicate that mouse and human CatSper are neither activated by cAMP directly nor indirectly via activation of PKA, and that activation of CatSper by membrane-permeable cAMP derivatives might represent a non-specific action of these compounds in sperm. The latter is supported by the finding that membrane-permeable derivatives of cGMP (e.g. 8-Br-cGMP) activate human CatSper only from the outside (Brenker et al., 2012). However, there are studies that contradict this concept and rather suggest a cAMP/PKA-activation of CatSper: in a study on human sperm, sizable bicarbonate-evoked Ca^2+^ signals were recorded (Spehr et al., 2004). Moreover, in a recent study by Orta et al (Orta et al., 2018), both 8-Br-cAMP and bicarbonate evoked Ca^2+^ influx via CatSper in mouse sperm. The Ca^2+^ influx by 8-Br-cAMP and bicarbonate was suppressed by inhibitors of PKA. Additionally, in patch-clamp recordings, CatSper-mediated membrane currents were enhanced by superfusion with bicarbonate and by cAMP in the pipette solution; inhibition of PKA suppressed the action of bicarbonate and intracellular cAMP. Based on these results, it was proposed that an increase of intracellular cAMP activates CatSper via activation of PKA (Orta et al., 2018).

Here, to solve this controversy, we studied the action of cAMP, membrane-permeable cAMP analogs, cAMP/PKA signaling, and PKA inhibitors in human sperm from healthy donors and patients that suffer from the deafness-infertility syndrome (DIS). In DIS patients, the *CATSPER2* gene is deleted (Avidan et al., 2003; Schiffer et al., 2020; Zhang et al., 2007), resulting in the loss of CatSper function (Brenker et al., 2018b; Schiffer et al., 2020; Smith et al., 2013). We demonstrate that human CatSper is neither activated by intracellular cAMP directly nor indirectly via activation of the cAMP/PKA-signaling pathway. In fact, membrane-permeable cAMP analogs and cAMP itself activate CatSper only from the outside via a so far unknown binding site that is distinct from that employed by steroids and prostaglandins. Furthermore, we found that several commonly used PKA inhibitors affect the activity of CatSper. The action of these drugs on CatSper does, however, not rest on the inhibition of PKA. Finally, we show that bicarbonate is prone to evoke artefactual alkaline-induced Ca^2+^ influx via CatSper that might be misinterpreted as cAMP/PKA-activation of the channel. Altogether, we conclude that the concept of a cAMP/PKA-activation of CatSper is rather based on chemical probes that are unspecific and, therefore, ill-suited to study the interplay of Ca^2+^- and cAMP-signaling pathways in sperm.

## Results

### CatSper is not activated by intracellular cAMP or cAMP/PKA signaling

To scrutinize whether CatSper is activated by intracellular cAMP or cAMP/PKA signaling, we studied the action of bicarbonate, IBMX, and adenosine in human sperm. Bicarbonate-activation of sAC rapidly increases cAMP (Fig. 1A) (Brenker et al., 2012) and activates PKA, i.e. cAMP levels and PKA activity peak within ≤ 60 s upon stimulation of sperm with bicarbonate (Battistone et al., 2013; Brenker et al., 2012). In sperm bathed in low concentrations of bicarbonate (e.g. 4 mM), isobutylmethylxanthine (IBMX) and adenosine evoke an increase of cAMP levels (Fig. 1A) (Brenker et al., 2012; Nolan et al., 2004; Schuh et al., 2006); IBMX and presumably also adenosine inhibit cAMP breakdown by phosphodiesterases (PDEs). However, neither bicarbonate nor IBMX or adenosine increased the intracellular Ca^2+^ concentration ([Ca^2+^]_i_) of human sperm, whereas CatSper activation by progesterone as a control evoked a biphasic Ca^2+^ response (Fig. 1B, C). Next, we recorded CatSper currents from human sperm by whole-cell patch clamping. In extracellular solution containing Ca^2+^ and Mg^2+^, we recorded only miniscule currents (Fig. 1D, E; HS); in Na^+^-based divalent-free solution, the prototypical monovalent CatSper currents were recorded (Fig. 1D, E; NaDVF) (Lishko et al., 2011; Strunker et al., 2011). The current amplitudes were similar in the absence and presence of cAMP in the pipette solution (Fig. 1D, E): mean CatSper inward currents at −80 mV were −20.7 ± 8.4 pA (n = 50) and −23.6 ± 11.9 pA (n = 16); mean outward currents at +80 mV were 53.3 ± 16.2 pA (n = 50) and 58.7 ± 21.4 pA (n = 16), respectively. Together, these results demonstrate that human CatSper is neither directly activated by intracellular cAMP nor indirectly by activation of the cAMP/PKA-signaling pathway (Brenker et al., 2012; Strunker et al., 2011). Of note, the intracellular cAMP level and activation of the cAMP/PKA-signaling pathways do not modulate stimulus-induced gating of human CatSper either: bicarbonate and IBMX did not affect CatSper activation by progesterone and simultaneous alkalization and depolarization (K8.6), respectively (Fig. 1H, I).

**Figure 1.**
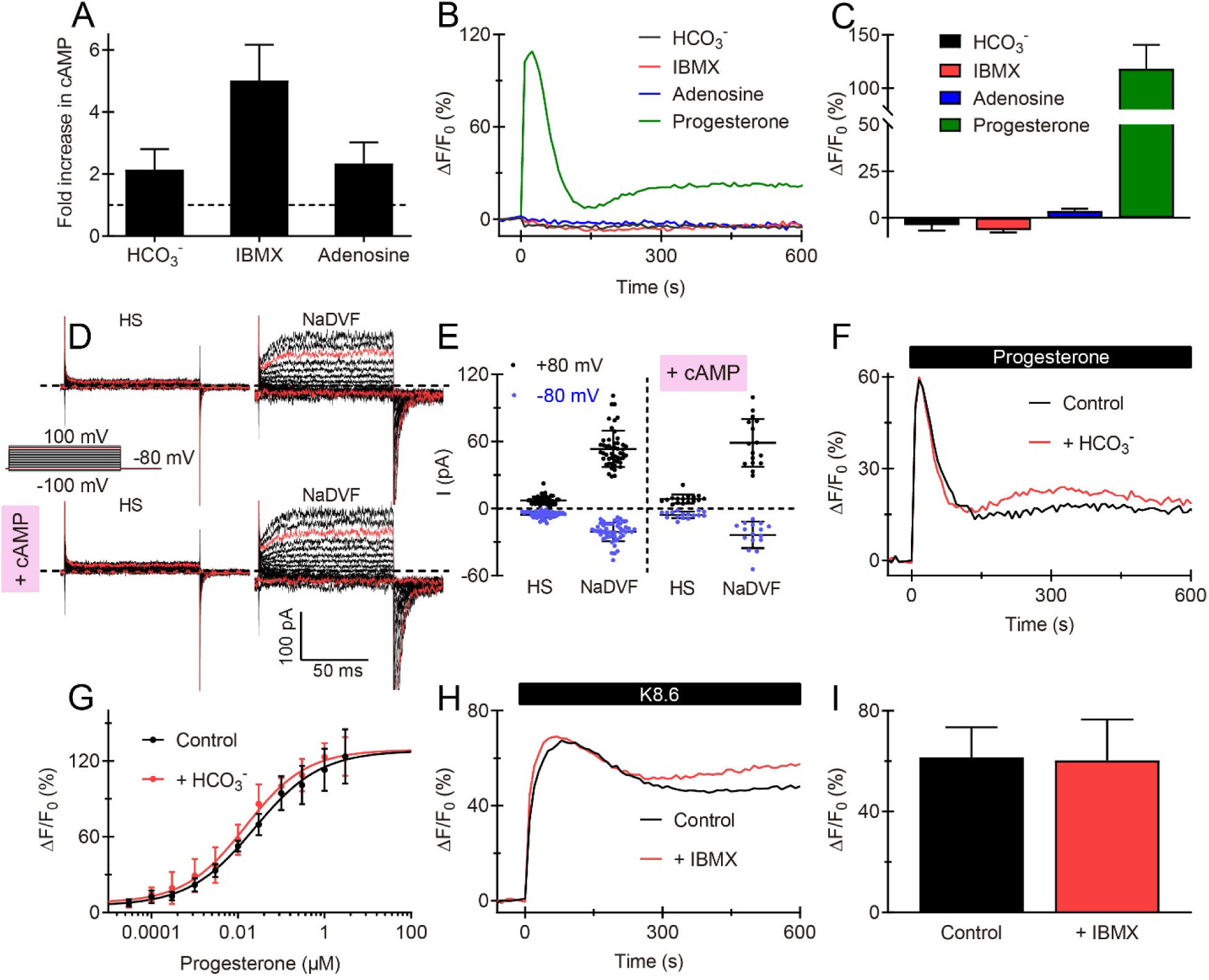
Human CatSper is not activated by intracellular cAMP. **(A)** Changes in total cAMP content in human sperm upon increasing bicarbonate from 0 mM to 25 mM HCO_3_^-^, or by IBMX (0.5 mM) or adenosine (0.1 mM) in the presence of 4 mM HCO_3_^-^. Dotted black line = 1 (control) (n = 3). **(B)** Representative Ca^2+^ signals evoked by increasing HCO_3_^-^ from 0 to 25 mM, or by application of IBMX (0.5 mM), adenosine (0.1 mM), or progesterone (10 μM) in the presence of 4 mM bicarbonate. [Ca^2+^]_i_ was monitored using a fluorescence plate reader (Strunker et al., 2011). Sperm were loaded with the fluorescent Ca^2+^ indicator Fluo-4. ΔF/F_0_ (%) indicates the percent change in fluorescence (ΔF) with respect to the mean basal fluorescence (F_0_) before application of stimulants at t = 0. **(C)** Mean amplitude of Ca^2+^ signals from B (n ≥ 3). **(D)** Representative whole-cell currents recorded from a human sperm cell at pH_i_ 7.3 in extracellular solution containing Mg^2+^ and Ca^2+^ (HS) and during perfusion with Na^+^-based divalent-free bath solution (NaDVF). Currents were evoked by stepping the membrane voltage from −100 to +100 mV (step 10 mV) from a holding potential of −80 mV in the absence (upper panel) or presence (lower panel) of cAMP (1 mM) in the pipette. Dotted black line: zero current level. Red traces: currents at +80 mV and −80 mV. **(E)** Steady-state current amplitudes at +80 mV and −80 mV in HS and NaDVF in the absence (n = 50, left) and presence of cAMP (1 mM) in the pipette (n = 16; right). **(F)** Ca^2+^ signals evoked by progesterone (10 nM) in sperm bathed in 0 (control) or 25 mM HCO_3_^-^. **(G)** Dose-response relationship for progesterone in the absence (control; EC_50_ = 23 ± 4 nM) and presence of HCO_3_^-^ (EC_50_ = 19 ± 19 nM) (n = 4). **(H)** Ca^2+^ signals in sperm bathed in 4 mM HCO_3_^-^ evoked by a simultaneous alkalization and depolarization (K8.6) in the absence (control) and presence of IBMX (0.5 mM). **(I)** Mean amplitude of Ca^2+^ signals from (H) (n = 4).

### Bicarbonate is prone to evoke alkaline-induced Ca^2+^ influx via CatSper

In some studies on mouse and human sperm, sizeable bicarbonate-evoked Ca^2+^ signals were recorded (Orta et al., 2018; Spehr et al., 2004). We entertained several eventualities that might explain these seemingly contradictory findings. We identified the pH of bicarbonate-containing buffers as a potential source of artifacts: the pH of buffers containing bicarbonate is tied to the partial pressure of CO_2_ (Kohn and Dunlap, 1998) and, therefore, unstable at ambient conditions. To illustrate this issue, we monitored the pH of HTF exposed to ambient air in the absence and presence of bicarbonate, using a fluorescent pH indicator. In the absence of bicarbonate, the pH remained stable at pH 7.35 (Fig. 2A). In the presence of bicarbonate, due to continuous degassing of CO_2_, the pH alkalized to pH 7.9 with a time constant (τ) of about 40 min (Fig. 2A). In sperm loaded with a fluorescent pH indicator, air-exposed (> 60 min) bicarbonate-HTF evoked an alkalization of the intracellular pH (pH_i_) that was similar to the pH_i_ alkalization evoked by alkaline (pH 7.8) HTF lacking bicarbonate (HTF7.8) or by the weak base ammonium chloride (NH_4_Cl) (Fig. 2B); stimulation of sperm with pH-controlled bicarbonate-HTF rather acidified pH_i_ (Fig. 2B). It is well-established that an intracellular alkalization by HTF7.8 or NH_4_Cl evokes Ca^2+^ influx via CatSper (Fig. 2C, D) (Rennhack et al., 2018; Schiffer et al., 2020; Strunker et al., 2011). Therefore, it is not surprising that air-exposed bicarbonate-HTF also evokes a sizeable [Ca^2+^]_i_ increase in human sperm (Fig. 2C, D). Thus, bicarbonate-evoked [Ca^2+^]_i_ increases observed in mouse and human sperm might reflect alkaline-rather than cAMP/PKA-induced CatSper activation, caused by unintended alkalization of bicarbonate buffers.

**Figure 2.**
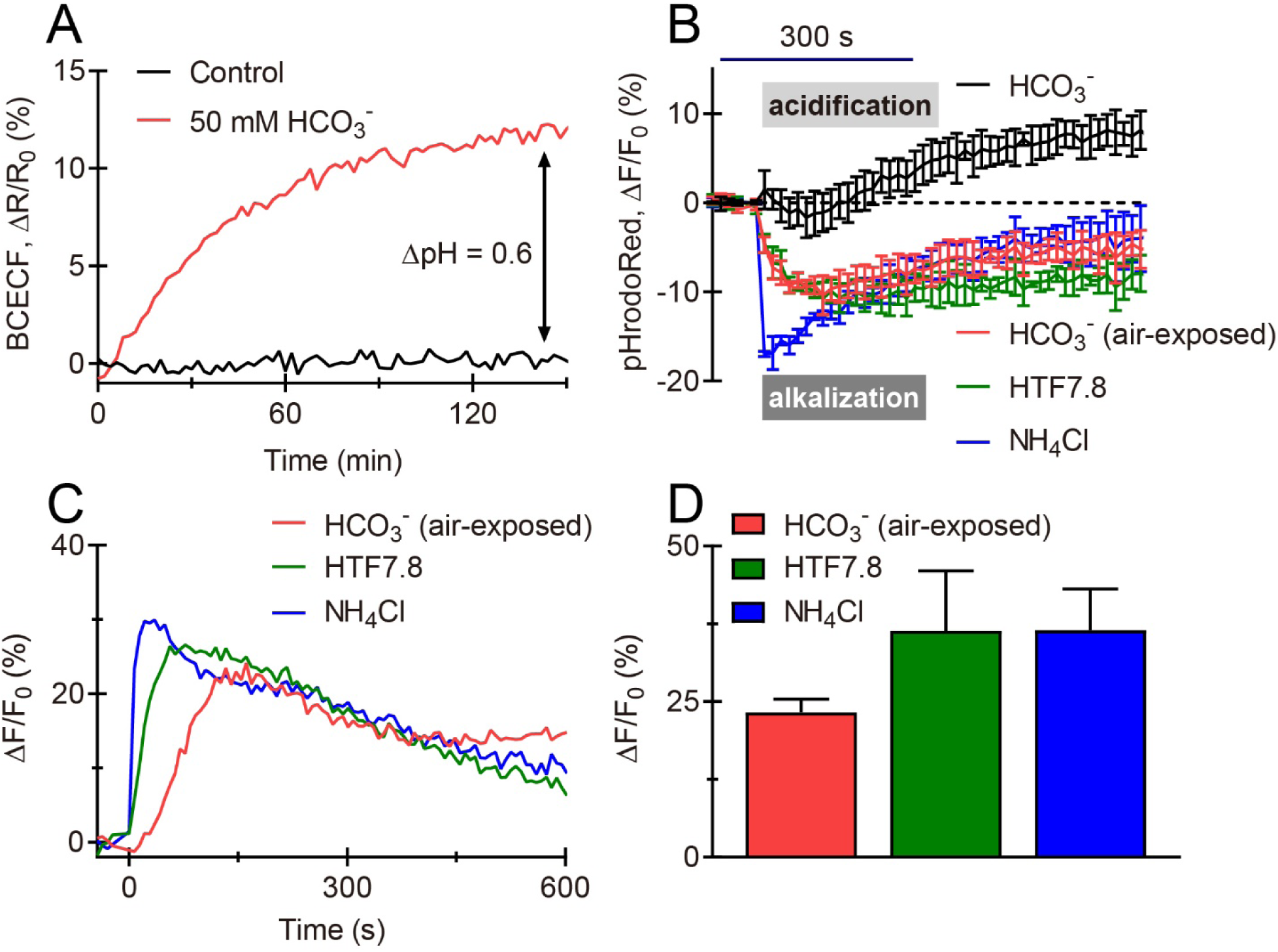
Bicarbonate is prone to evoke alkaline-induced Ca^2+^ influx via CatSper. **(A)** Changes in the pH of HTF buffer in the absence (control) and presence of HCO_3_^-^ (50 mM) at ambient air, measured with the fluorescent pH-indicator BCECF. **(B)** pH_i_ changes evoked by pH-controlled HTF containing HCO_3_^-^ (25 mM), air-exposed, alkaline HTF containing HCO_3_^-^ (25 mM), alkaline HCO_3_^-^-free HTF (pH 7.8, HTF7.8), and NH_4_Cl (5 mM) in human sperm loaded with the fluorescent pH indicator pHrodoRed. Dotted black line indicates Y = 0 (n = 3). **(C)** Ca^2+^ signals evoked by air-exposed, alkaline HTF containing HCO_3_^-^ (25 mM), HTF7.8, and NH_4_Cl (5 mM). **(D)** Mean amplitude of Ca^2+^ signals from (C) (n = 3).

### Membrane-permeable cAMP analogs activate CatSper only from the outside

Next, considering that human CatSper is not activated by intracellular cAMP, we set out to elucidate the mechanism underlying the Ca^2+^ influx evoked by membrane-permeable cAMP analogs. To this end we studied the action of the most common cAMP analogs used to mimic the action of intracellular cAMP (8-CPT-cAMP, 8-Br-cAMP, Db-cAMP) or to activate PKA (6-Bnz-cAMP) or EPAC (8-pCPT-2-O-Me-cAMP-AM, i.e. 007-AM) (Schmidt et al., 2013). Any of these cAMP analogs evoked a biphasic [Ca^2+^]_i_ increase, and the amplitude of the Ca^2+^ response rose in a dose-dependent fashion (Fig. 3A, B; Supplementary Fig. 1A). The potency of the cAMP analogs varied: 8-pCPT-2-O-Me-cAMP-AM and 8-CPT-cAMP increased [Ca^2+^]_i_ already at micromolar concentrations, whereas the action of 8-Br-cAMP, Db-cAMP, and 6-Bnz-cAMP commenced only at ≥ 1 mM (Fig. 3B; Supplementary Fig. 1A). Because we used the Na^+^ salts of the cAMP analogs, we studied the action of NaCl as a surrogate for the vehicle. Only at > 10 mM, the vehicle evoked a small Ca^2+^ response on its own (Fig. 3B). To scrutinize whether the [Ca^2+^]_i_ increase evoked by cAMP analogs is mediated by CatSper, we studied their action in CatSper-deficient sperm from infertile patients that lack the *CATSPER2* gene (Brenker et al., 2018b; Schiffer et al., 2020). In CatSper-deficient sperm, the Ca^2+^ response evoked by 8-CPT-cAMP, 8-Br-cAMP, 6-Bnz-cAMP, and 8-pCPT-2-O-Me-cAMP-AM was abolished, and Db-cAMP evoked only a small residual Ca^2+^ signal (Fig. 3A; Supplementary Fig. 1A). Furthermore, in whole-cell patch-clamp recordings, superfusion of sperm with 8-CPT-cAMP enhanced monovalent currents at −80 mV and +80 mV by 2.47 ± 0.8-fold and by 1.22 ± 0.16-fold (n = 8), respectively, confirming that 8-CPT-cAMP activates CatSper. The vehicle had only a minuscule, if any, action on the amplitude of CatSper currents (Supplementary Fig. 1B-D); and in CatSper-deficient sperm, 8-CPT-cAMP did not affect residual monovalent currents. (Supplementary Fig. 2A, B). These results demonstrate that CatSper is promiscuously activated by structurally and functionally diverse membrane-permeable analogs of cAMP. Of note, CatSper is not only activated by cAMP analogs, but also by analogs of cGMP such as 8-CPT-cGMP or 8-Br-cGMP (Supplementary Fig. 3A, B) (Brenker et al., 2012), which activate human CatSper only via an extracellular binding site (Brenker et al., 2012). We examined whether cAMP analogs activate CatSper by a similar mechanism. To this end, we first tested by Ca^2+^ fluorimetry whether CatSper might also be activated by extracellular application of cAMP itself, which hardly permeates the cell membrane. Indeed, not only its membrane-permeable analogs, but also cAMP (at ≥ 1 mM), evoked a [Ca^2+^]_i_ increase in human sperm (Fig. 4A, B). The cAMP-evoked Ca^2+^ response was abolished in sperm that lack functional CatSper (Fig. 4A). Next, we examined the kinetics of the Ca^2+^ signals evoked by 8-CPT-cAMP in a stopped-flow apparatus. Upon rapid mixing of sperm with 8-CPT-cAMP, [Ca^2+^]_i_ rose with no measurable latency within the time resolution of the system (36 ms) (Fig. 4C, D). Finally, in patch-clamp recordings, superfusion of sperm with cAMP enhanced monovalent CatSper currents at −80 mV and +80 mV by 1.58 ± 0.17-fold and by 1.36 ± 0.4-fold (n = 4) (Fig. 5A-C), respectively. The presence of cAMP or 8-CPT-cAMP in the pipette solution did not suppress activation of CatSper by superfusion with 8-CPT-cAMP (Fig. 5D-I): 8-CPT-cAMP enhanced inward and outward currents by 1.88 ± 0.45-fold (−80 mV) and 1.03 ± 0.13-fold (+80 mV) (n = 8) in the presence of cAMP (n = 8); and by 1.86 ± 0.2-fold (−80 mV) and 1.04 ± 0.08-fold (+80 mV) in the presence of 8-CPT-cAMP (n = 4). Altogether, the activation of CatSper by extracellular cAMP, the failure of intracellular cAMP or 8-CPT-cAMP to suppress CatSper activation by superfusion with 8-CPT-cAMP, and the virtually instantaneous onset of 8-CPT-cAMP-evoked Ca^2+^ responses indicate that cAMP and membrane-permeable cAMP analogs activate CatSper only from the extracellular space.

**Figure 3.**
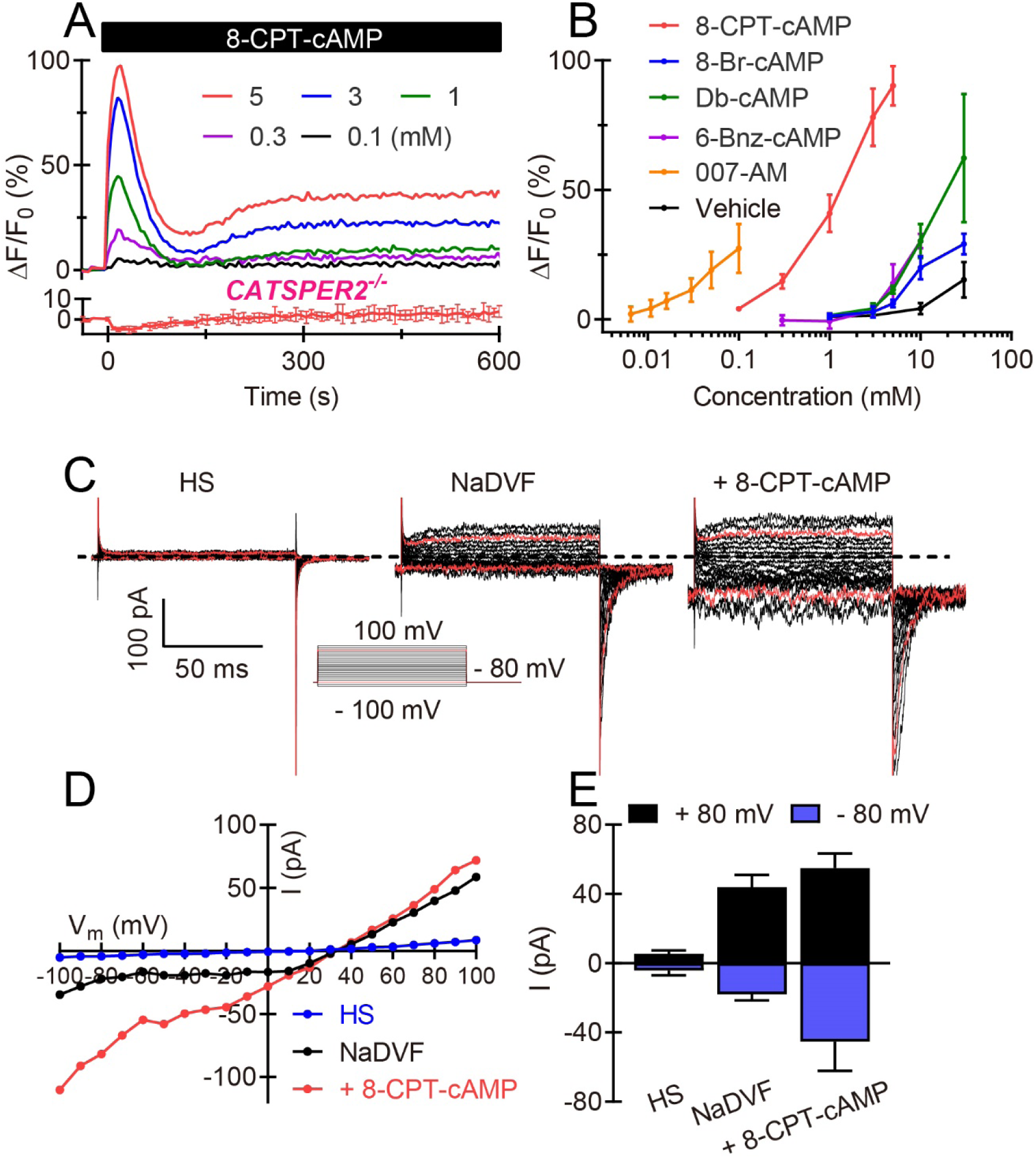
Membrane-permeable cAMP analogs activate CatSper in human sperm. **(A)** Representative Ca^2+^ signals evoked by 8-CPT-cAMP in sperm from a healthy donor (upper panel), and averaged Ca^2+^ signal (lower panel, n = 3) in sperm lacking functional CatSper channels (*CATSPER2^-/-^*). **(B)** Mean amplitudes of Ca^2+^ signals evoked by membrane-permeable cAMP analogs and by the vehicle (NaCl) (n = 3). **(C)** Representative whole-cell currents recorded from human sperm at pH_i_ 7.3 in extracellular solution containing Ca^2+^ and Mg^2+^ (HS), in divalent-free Na^+^-based bath solution (NaDVF), and during perfusion with NaDVF containing 8-CPT-cAMP (5 mM). Dotted black line: zero current level. Red traces: currents at +80 mV and −80 mV. **(D)** Steady-state current-voltage relationship from (C). **(E)** Mean current amplitudes at +80 mV and −80 mV in HS, in NaDVF, and in NaDVF containing 5 mM 8-CPT-cAMP (n = 8).

**Figure 4.**
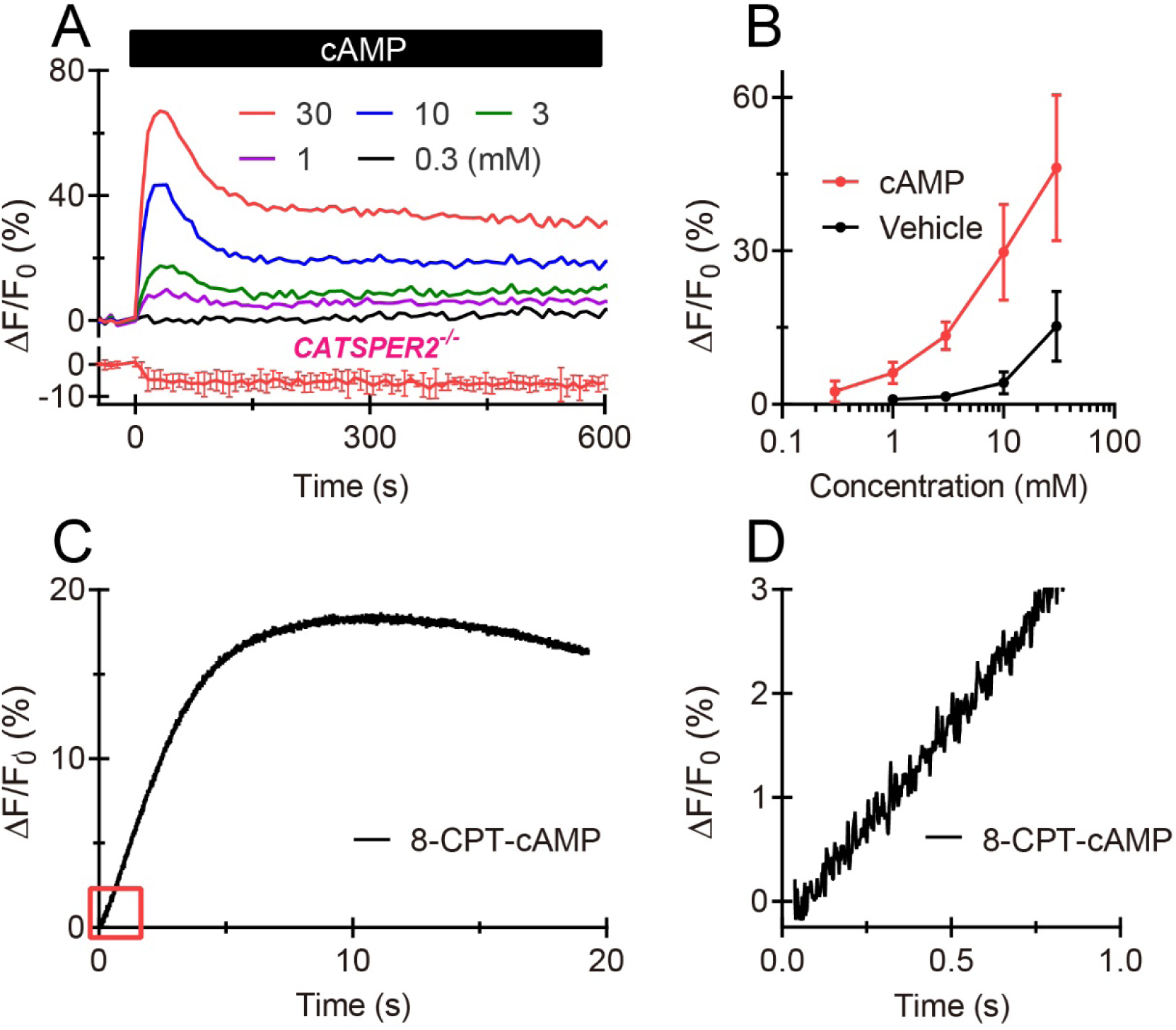
Activation of CatSper by extracellular cAMP and kinetics of the 8-CPT-cAMP-evoked Ca^2+^ response in human sperm. **(A)** Representative Ca^2+^ signals evoked by cAMP in sperm from a healthy donor (upper panel), and averaged Ca^2+^ signal (lower panel, n = 3) in sperm lacking functional CatSper channels (*CATSPER2^-/-^*). **(B)** Mean amplitudes of Ca^2+^ signals evoked by cAMP (n = 4) and the vehicle (NaCl) (n = 3). **(C)** Ca^2+^ signal evoked by rapid mixing of sperm with 8-CPT-cAMP (5 mM) in a stopped-flow apparatus. **(D)** Onset of the 8-CPT-cAMP-evoked Ca^2+^ signal from (C) shown on an extended time scale.

**Figure 5.**
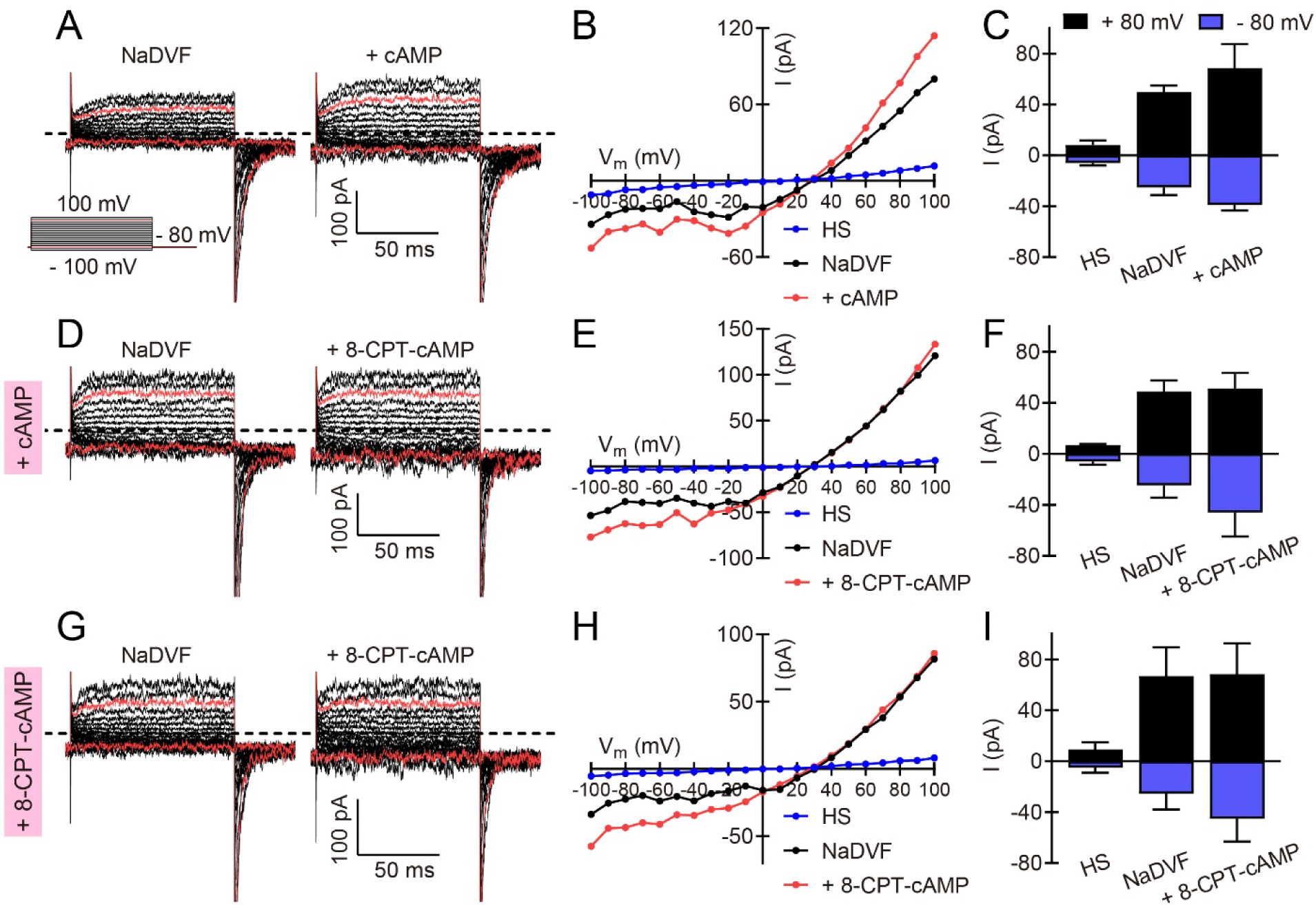
cAMP and 8-CPT-cAMP activate human CatSper from the outside. **(A, D, and G)** Representative whole-cell currents at pH_i_ 7.3 in divalent-free Na^+^-based bath solution (NaDVF) and in NaDVF containing 10 mM cAMP (A) or 5 mM 8-CPT-cAMP (D and G). Currents were evoked in the absence (A) or in the presence of 1 mM cAMP (D) or 5 mM 8-CPT-cAMP (G) in the pipette. Dotted black line: zero current level. Red traces: currents at +80 mV and −80 mV. **(B, E, and H)** Steady-state current-voltage relationships from (A), (D), and (G), respectively. **(C, F, and I)** Mean current amplitudes at +80 mV and −80 mV in HS, in NaDVF, and in NaVDF containing 10 mM cAMP (C, n = 4) or 5 mM 8-CPT-cAMP (F, n = 8; I, n = 4). Currents were evoked in the absence (C) or presence of 1 mM cAMP (F) or 5 mM 8-CPT-cAMP (I) in the pipette.

### Human CatSper is activated by cyclic nucleotides via a so far unknown binding site

We wondered whether cyclic nucleotides compete with steroids and/or prostaglandins to activate CatSper. Ca^2+^-fluorimetric cross-desensitization experiments revealed that steroids and prostaglandins employ different binding sites to activate human CatSper (Brenker et al., 2018b; McBrinn et al., 2019; Schaefer et al., 1998; Strunker et al., 2011). We used this approach to study the mechanism of cyclic nucleotide-induced CatSper activation. Validating our experimental conditions, pre-incubation of sperm with a saturating concentration of progesterone abolished the Ca^2+^ response evoked by 17-OH-progesterone, but not that evoked by PGE_1_ (Fig. 6A, B); and pre-incubation with a saturating concentration of PGE_1_ abolished the Ca^2+^ response evoked by PGE_2_, but not that evoked by progesterone (Fig. 6C, D). Pre-incubation with saturating concentration of progesterone or PGE_1_, however, did not abolish Ca^2+^ responses evoked by cAMP, cAMP analogs, or cGMP analogs (Fig. 6A-D); vice versa, pre-incubation of sperm with 8-CPT-cAMP or 8-CPT-cGMP did not abolish progesterone- or PGE_1_-evoked Ca^2+^ responses (Fig. 6E-H). The lack of cross-desensitization demonstrates that cyclic nucleotides do not compete with progesterone or prostaglandins to activate CatSper. However, pre-incubation with 8-CPT-cAMP or 8-CPT-cGMP abolished the responses evoked by cAMP as well as by other analogs of cAMP or cGMP (Fig. 6E-G), indicating that cAMP, cAMP analogs, and cGMP analogs compete with each other to activate CatSper. Thus, cyclic nucleotides, steroids, and prostaglandins activate human CatSper via three distinct binding sites.

**Figure 6.**
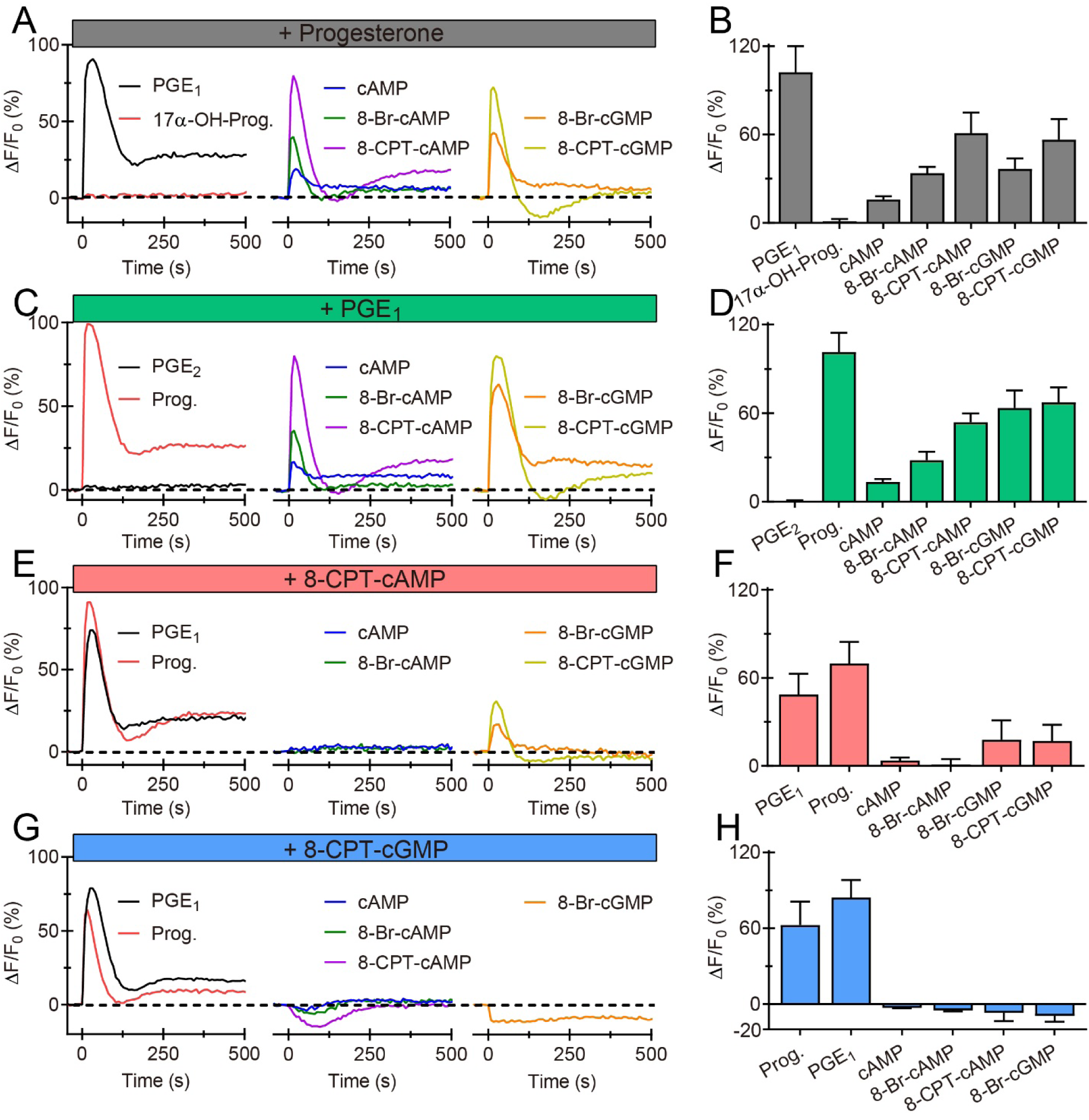
Cyclic nucleotides, steroids, and prostaglandins activate human CatSper via three distinct binding sites. **(A, B)** Representative Ca^2+^ signals (A) and mean signal amplitudes (B; n ≥ 3) evoked by 17α-OH-Progesterone (2 μM), PGE_1_ (2 μM), cAMP (10 mM), 8-Br-cAMP (10 mM), 8-CPT-cAMP (5 mM), 8-Br-cGMP (10 mM), and 8-CPT-cGMP (5 mM) in sperm bathed in progesterone (10 µM). **(C, D)** Representative Ca^2+^ signals and mean signal amplitudes (n ≥ 3) evoked by PGE_2_ (2 µM), progesterone (2 µM), and the indicated cyclic nucleotides in sperm bathed in PGE_1_ (2 µM). **(E, F)** Representative Ca^2+^ signals and mean signal amplitudes (n ≥ 3) evoked by PGE_1_, progesterone, and the indicated cyclic nucleotides in sperm bathed in 8-CPT-cAMP (5 mM), **(G, H)** Representative Ca^2+^ signals and mean signal amplitudes (n ≥ 3) evoked by PGE_1_, progesterone, and the indicated cyclic nucleotides in sperm bathed in 8-CPT-cGMP (5 mM).

### The action of PKA inhibitors on CatSper does not rest on inhibition of PKA

Given that CatSper is not activated by cAMP/PKA signaling, how can inhibitors of PKA, i.e. H89 and PKI 14-22, suppress CatSper-mediated Ca^2+^ influx in human and mouse sperm (Baron et al., 2016; Orta et al., 2018). To solve this conundrum, we studied the action of the five most common pharmacological probes (i.e. PKI 5-24, Rp-cAMPS, KT 5720, H89, and PK 14-22) in human sperm. The action of these drugs was heterogeneous: PKI 5-24 and Rp-cAMPS affected neither resting Ca^2+^ levels nor CatSper-mediated Ca^2+^ influx evoked by 8-CPT-cAMP, progesterone, PGE_1_, or alkalization (NH_4_Cl) (Fig. 7A-D). Similarly, KT 5720 did not suppress ligand- or alkaline-induced Ca^2+^ influx via CatSper (Fig. 7A-D). However, on its own, KT 5720 evoked a small, transient [Ca^2+^]_i_ increase that was abolished in CatSper-deficient human sperm (Fig. 7A, B; Supplementary Fig. 4A). In patch-clamp recordings from human sperm, KT 5720 increased the amplitude of monovalent CatSper inward (−80 mV) and outward (+80 mV) currents by 1.48 ± 0.47-fold and 1.33 ± 0.11-fold, respectively (Fig. 7E, F, n = 5). The action of H89 was particularly complex: H89 on its own evoked a transient Ca^2+^ increase, followed by a [Ca^2+^]_i_ decrease below resting levels (Fig. 8A, B). The dose-response relation for the Ca^2+^ transient was bell shaped, i.e. the signal amplitude grew with increasing concentrations, saturated at about 10 µM, and decreased again at >10 µM (Fig. 8B). The H89-evoked Ca^2+^ response was abolished in CatSper-deficient human sperm, indicating that H89 acts via CatSper (Supplementary Fig. 4A). Of note, in the presence of H89, both ligand- and alkaline-evoked Ca^2+^ influx via CatSper was strongly attenuated (Fig. 8C, D); and in patch-clamp recordings, H89 reversibly inhibited monovalent CatSper currents (Fig. 8E, F; Supplementary Fig. 4B). Finally, PKI 14-22 evoked a small decrease of [Ca^2+^]_i_ on its own, which was abolished in CatSper-deficient sperm (Fig. 8A; Supplementary Fig. 4A). PKI 14-22 strongly attenuated alkaline-evoked Ca^2+^ influx via CatSper, whereas ligand-evoked Ca^2+^ influx was only slightly suppressed (Fig. 8C, D). In patch-clamp recordings, PKI 14-22 inhibited monovalent CatSper currents (Supplementary Fig. 4C).

**Figure 7.**
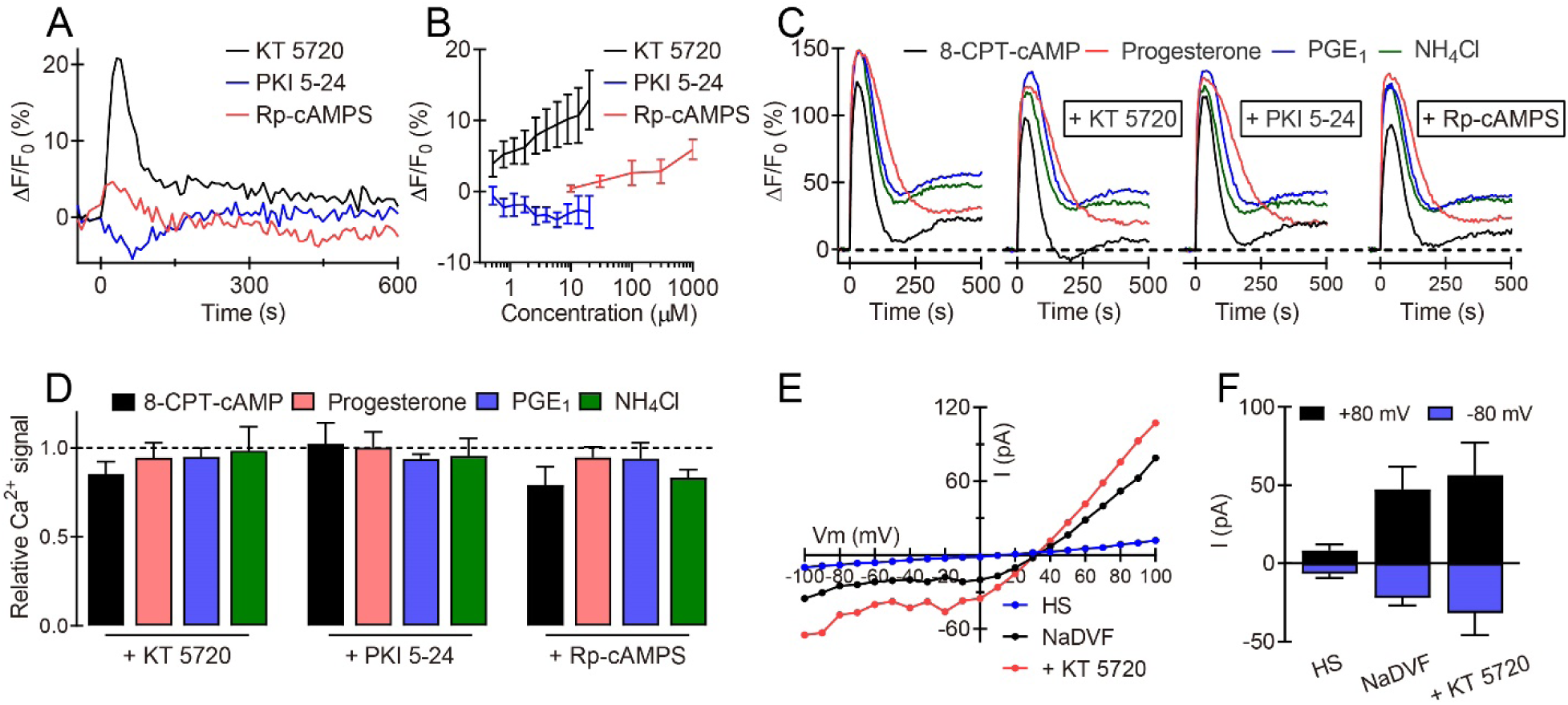
Action of PKA inhibitors on CatSper in human sperm. **(A)** Representative Ca^2+^ signals evoked by KT 5720 (20 μM), PKI 5-24 (20 μM), and Rp-cAMPS (1 mM). **(B)** Mean amplitudes of Ca^2+^ signals evoked by KT 5720, PKI 5-24, and Rp-cAMPS (n ≥ 3). **(C)** Ca^2+^ signals evoked by 8-CPT-cAMP (5 mM), progesterone (10 μM), PGE_1_ (10 μM), and NH_4_Cl (10 mM), in the absence and presence of KT 5720 (20 μM), PKI 5-24 (20 μM), or Rp-cAMPS (1 mM). **(D)** Mean amplitude of Ca^2+^ signals in the presence of KT 5720 (20 μM), PKI 5-24 (20 μM), or Rp-cAMPS (1 mM) relative to the amplitude evoked in the absence of any drug (n ≥ 4). **(E)** Representative steady-state current-voltage relationship of currents recorded at pH_i_ of 7.3 in extracellular solution containing Ca^2+^ and Mg^2+^ (HS), in divalent-free Na^+^-based bath solution (NaDVF), and in NaDVF containing KT 5720 (20 μM). **(F)** Mean current amplitudes at +80 mV and −80 mV recorded in HS, in NaDVF, and in NaDVF containing 20 μM KT 5720 (n = 5).

**Figure 8.**
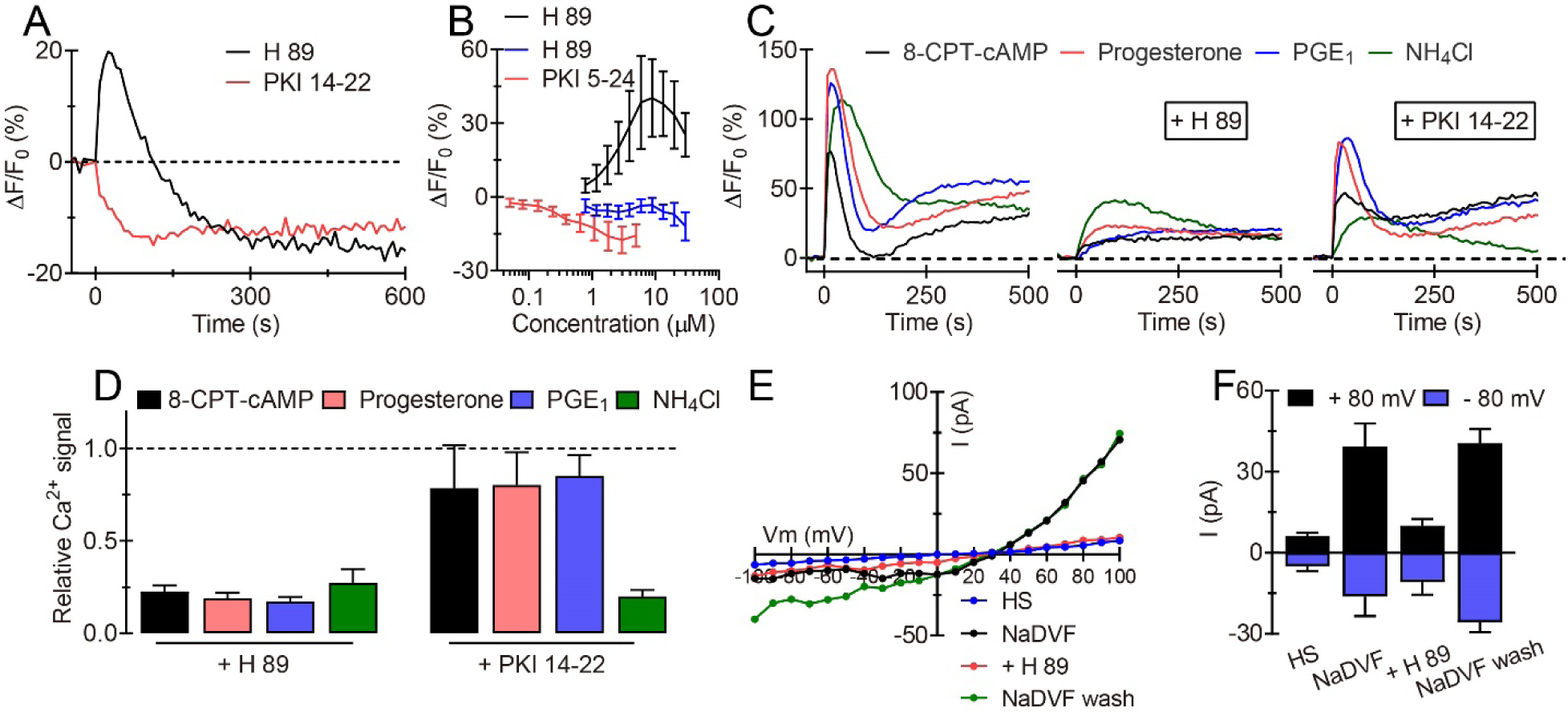
The PKA inhibitors H89 and PKI 14-22 inhibit human CatSper. **(A)** Representative Ca^2+^ signals evoked by H89 (30 μM) and PKI 14-22 (5 μM). **(B)** Mean amplitude of Ca^2+^ signals evoked by H89 (black, peak phase; blue, sustained phase) and PKI 14-22 (n ≥ 6). **(C)** Ca^2+^ signals evoked by 8-CPT-cAMP (5 mM), progesterone (10 μM), PGE_1_ (10 μM), and NH_4_Cl (10 mM) in the absence and presence of H89 (30 μM) or PKI 14-22 (5 μM). **(D)** Mean amplitude of Ca^2+^ signals in the presence of H89 or PKI 14-22 relative to the amplitude evoked in the absence of any drug (n ≥ 4). **(E)** Representative steady-state current-voltage relationship of currents recorded at pH_i_ 7.3 in extracellular solution containing Ca^2+^ and Mg^2+^ (HS), in divalent-free Na^+^-based bath solution (NaDVF), and in NaDVF containing H89 (30 μM). **(F)** Mean current amplitudes at +80 mV and −80 mV in HS, in NaDVF, and in NaDVF containing H89 (n ≥ 3).

In summary, H89 and PKI 14-22 indeed inhibit CatSper. This action rests, however, on a direct inhibition of CatSper itself rather than on inhibition of PKA: in patch-clamp recordings, both H89 and PKI 14-22 suppress basal CatSper currents recorded under conditions that do not foster PKA signaling. Moreover, H89 and PKI 14-22 suppress alkaline-evoked Ca^2+^ influx via CatSper, which does not involve activation of PKA. Thus, H89 and PKI 14-22 are not suited to study the interconnection of cAMP, PKA, and Ca^2+^ signaling in human sperm. In contrast to H89 and PKI 14-22, KT 5720 rather activates the channel, which renders also KT 5720 ill-suited for studies of PKA and/or CatSper function in sperm. Of note, the nonspecific action of the PKA inhibitors on CatSper does not come as a surprise. In fact, these drugs are well known to be rather nonspecific (Murray, 2008).

## Discussion

Our results reinforce the conclusion that human CatSper is not activated by intracellular cAMP or cAMP/PKA signaling (Brenker et al., 2012). In fact, the concept of a cAMP/PKA-activation of CatSper rests on pharmacological tools that are, across the board, prone to unspecific effects. First, if not properly controlled for pH, bicarbonate solutions can activate CatSper via an increase of pH_i_. Second, the common PKA inhibitors H89 and PKI 14-22 directly block CatSper, whereas KT 5720 activates the channel. Finally, membrane-permeable analogs of cyclic nucleotides activate CatSper only via an extracellular binding site. These findings enlarge the list of drugs that are frequently used to interfere with sperm signaling, but directly affect CatSper. For example, the popular PLC and PDE inhibitors U73122 and trequensin, respectively, as well as a diverse array of synthetic chemicals evoke Ca^2+^ influx via CatSper (Brenker et al., 2012; Diao et al., 2017; McBrinn et al., 2019; Schiffer et al., 2014; Tavares et al., 2013). By contrast, MDL12330A, an inhibitor of transmembrane ACs, and RU1968, a sigma-receptor agonist, both block CatSper (Brenker et al., 2012; Rennhack et al., 2018; Schiffer et al., 2014). Moreover, also inhibitors of EPAC, another cAMP effector that reportedly controls [Ca^2+^]_i_ in sperm (Itzhakov et al., 2019; Lucchesi et al., 2016), directly act on CatSper. The EPAC inhibitors CE3F4, ESI 05, and HJC 0350 activate human CatSper, whereas ESI 09 blocks the channel (Supplementary Fig. 5). Thus, the most common EPAC inhibitors are also not suited to investigate the role of EPAC in Ca^2+^-signaling of sperm. Altogether, these results caution against rash interpretation in mechanistic terms of results derived from experiments with pharmacological tools that seemingly control [Ca^2+^]_i_ and/or CatSper. Because of the miniscule flagellar volume, even minute changes in CatSper activity strongly affect [Ca^2+^]_i_ and, thereby, downstream signaling events. Therefore, quite general, using pharmacological tools to study signaling in sperm requires careful assessment of potential off-target actions on CatSper.

We did not examine whether the off-target action of membrane-permeable cAMP/cGMP analogs and PKA inhibitors on CatSper are similar in human and mouse sperm. Examples of species-specific ligand actions on CatSper are steroids and prostaglandins that activate CatSper in human (Brenker et al., 2012; Lishko et al., 2011; Strunker et al., 2011), but not in mouse sperm (Tamburrino et al., 2015). Therefore, we cannot exclude that there are differences concerning the cAMP/PKA-activation of CatSper among species. However, H89 suppressed alkaline-induced membrane currents also in mouse sperm (Orta et al., 2018), suggesting that H89 also directly blocks mouse CatSper. Moreover, a large number of independent studies by different groups consistently provided no evidence that a rise of cAMP and ensuing PKA activation activates mouse CatSper (Carlson et al., 2007; Carlson et al., 2003; Jansen et al., 2015; Nolan et al., 2004; Schuh et al., 2006; Wennemuth et al., 2003a). Together, this comprehensive body of evidence strongly suggests that also mouse CatSper is not directly activated by intracellular cAMP/PKA signaling.

Of note, the activity of CatSper is remodeled during the capacitation process: the voltage dependence of CatSper shifts to more negative potentials (Lishko et al., 2011), and human CatSper becomes more sensitive to progesterone and prostaglandins (Bedu-Addo et al., 2005; Strunker et al., 2011). The underlying mechanism(s) are unknown, but might involve capacitation-associated remodeling of the membrane-lipid environment (Kawai et al., 2019), intracellular pH (Puga Molina et al., 2018), and/or V_m_ (Puga Molina et al., 2018) rather than cAMP signaling.

In the past decade, a picture has emerged that human CatSper serves as a polymodal chemosensor that translates the chemical code of the oviductal environment to changes in [Ca^2+^]_i_ and motility (Alasmari et al., 2013; Brenker et al., 2012; Brenker et al., 2018b; Diao et al., 2014; Diao et al., 2017; Lishko et al., 2011; Strunker et al., 2011; Williams et al., 2015). Oviductal steroids and prostaglandins activate CatSper in a highly synergistic fashion (Brenker et al., 2018a) via two distinct binging sites (Lishko et al., 2011; McBrinn et al., 2019; Schaefer et al., 1998; Strunker et al., 2011). Progesterone has been proposed to activate human CatSper via the receptor alpha/beta hydrolase domain-containing protein 2 (ABHD2) (Miller et al., 2016). By contrast, the mechanism of CatSper activation by prostaglandins is elusive, except that it does not involve ABHD2, classical G protein-coupled receptors, and second messengers (Brenker et al., 2012; Brenker et al., 2018c; Lishko et al., 2011; McBrinn et al., 2019; Miller et al., 2016; Schaefer et al., 1998; Strunker et al., 2011). Our results demonstrate that CatSper is also controlled by an extracellular binding site that accommodates neither steroids nor prostaglandins, but cyclic nucleotides. Yet, extracellular cAMP activates CatSper only at concentrations ≥ 1 mM, which exceeds physiological extracellular cAMP concentrations by several orders of magnitude. This indicates that the true physiological ligand of the cyclic nucleotide-binding site is to be deorphanized. Moreover, future studies are required to elucidate the molecular identity of this novel binding site, and it needs to be examined whether activation of CatSper via this third binding site evokes motility responses that are similar or distinct to that evoked by steroids or prostaglandins.

## Materials and Methods

### Reagents

cAMP, cGMP, and their derivatives (sodium salts) were obtained from BIOLOG Life Science Institute (Bremen, Germany). 8-pCPT-2-O-Me-cAMP-AM, H89 dihydrochloride, PKI 14-22 amide myristoylated, ESI 09, CE3F4, HJC 0350, and ESI 05 were from Tocris (Minnesota, USA). Prostaglandin E_1_ (PGE_1_), Prostaglandin E_2_ (PGE_2_), and PKI 5-24 were obtained from Cayman (Hamburg, Germany). Fluo-4-AM, pHrodo Red-AM, and BCECF were obtained from Invitrogen (California, USA). Human serum albumin (HSA) was obtained from Irvine Scientific (Santa Ana, USA). All other chemicals were from Sigma-Aldrich.

### Human sperm preparation

Semen samples were obtained from volunteers and DIS patients with prior written consent, under approval from the ethical committees of the medical association Westfalen-Lippe and the medical faculty of the University of Münster (4INie). Semen samples were produced by masturbation and ejaculated into plastic containers. The samples were allowed to liquefy for 15∼30 min at 37°C and motile sperm were purified by a swim-up procedure: liquefied semen (0.5–1 ml) was directly layered in a 50 ml falcon tube below 4 ml of human tubal fluid (HTF) medium, containing (in mM): 97.8 NaCl, 4.69 KCl, 0.2 MgSO_4_, 0.37 KH_2_PO_4_, 2.04 CaCl_2_, 0.33 Na-pyruvate, 21.4 lactic acid, 4 NaHCO_3_, 2.78 glucose, and 21 HEPES, pH 7.35 (adjusted with NaOH). Alternatively, the liquefied semen was diluted 1:10 with HTF, and sperm, somatic cells, and cell debris were pelleted by centrifugation at 700g for 20 min (37°C). The pellet was resuspended in the same volume HTF, 50 ml falcon tubes were filled with 5 ml of the suspension, and cells were pelleted by centrifugation at 700 g for 5 min (37°C). In both cases, motile sperm were allowed to swim up into HTF for 60–90 min at 37 °C. Sperm were washed two times (700g, 20 min, 37°C) and re-suspended in HTF containing 3 mg/ml HSA at a density of 1 × 10^7^ sperm/ml for measurement of changes in intracellular Ca^2+^, pH, or cAMP as well as for patch-clamp recordings. To study the action of bicarbonate, sperm were purified by swim-up in HTF medium lacking NaHCO_3_, which was substituted with NaCl. Sperm were washed two times (700g, 20 min, 37°C) and re-suspended in bicarbonate-free HTF containing 3 mg/ml HSA.

### Measurement of changes in [Ca^2+^]_i_ and pH

Changes in [Ca^2+^]_i_ and pH_i_ were measured in sperm loaded with the fluorescent Ca^2+^ and pH indicator, Fluo-4-AM and pHrodo Red-AM, respectively, at 30°C in 384 multi-well plates in a fluorescence plate reader (Fluostar Omega, BMG Labtech, Ortenberg, Germany) (Strunker et al., 2011). Sperm were loaded with Fluo-4-AM (5 μM, 20 min) and pHrodo Red-AM (5 μM, 30 min) at 37°C in the presence of Pluronic F-127 (0.05% w/v). After incubation, excess dye was removed by centrifugation (700g, 5 min, room temperature). Sperm were resuspended in HTF at a concentration of 5 × 10^6^/ml. The wells were filled with 50 μl of the sperm suspension and the fluorescence was excited at 480 nm (Fluo-4, pHrodo Red) and fluorescence emission was recorded at 520 nm. Changes in Fluo-4 and pHrodo Red fluorescence are depicted as ΔF/F_0_ (%), that is, the change in fluorescence (ΔF) relative to the mean basal fluorescence (F_0_) before application of buffer or stimuli (25 µl), to correct for intra- and inter-experimental variations in basal fluorescence among individual wells.

To study the action of bicarbonate, wells were filled with 40 µl of the sperm suspension in bicarbonate-free HTF and 40 µl of HTF containing 50 mM bicarbonate were injected. Of note, the bicarbonate-HTF was stored in air-tight tubes filled to capacity to avoid alkalization due to exposure to room air. Moreover, right prior to the experiments, the pH was checked again and, if required, (re)adjusted. To measure Ca^2+^ signals evoked by simultaneous alkalization/depolarization, wells were filled with 40 µl of the sperm suspension and 40 µl of K8.6 solution, containing (in mM): 98.5 KCl, 0.2 MgSO_4_, 0.37 KH_2_PO_4_, 2.04 CaCl_2_, 21.4 lactic acid, 4 KHCO_3_, 2.78 glucose, and 21 HEPES, pH 9.3 (adjusted with KOH), was injected; the final pH in the well was 8.6. To measure the pH change of HTF containing 50 mM NaHCO_3_ exposed to room air, 5 μM BCECF was used. The wells were filled with 50 µl HTF and BCECF was exited at 440 and 480 nm (dual excitation, BCECF) and the emission was recorded at 520 nm over 2 hours. Changes in BCECF-fluorescence ratio (R, 480/440 nm) are depicted as ΔR/R (%), that is, the change in ratio (ΔR) relative to the mean ratio (R0) measured within the first minute of the recording.

Stopped-flow experiments were performed as described before (Brenker et al., 2012; Strunker et al., 2011) with some modifications (see also (Hamzeh et al., 2019)). Briefly, In a SFM-400 stopped-flow device (Bio-Logic, France), a suspension of Fluo-4-loaded sperm (1 × 10^7^/ml) in HTF was rapidly mixed (1:1; flow rate = 1 ml/s) with HTF containing 10 mM 8-CPT-cAMP. Fluorescence was excited with a blue light-emitting diode (LED; M470D2, Thorlabs, Germany; powered with a custom-made power supply) that was modulated at 10 kHz using a function generator (4060MV, PeakTech, Germany). The light was passed through a 475/28 nm excitation filter (Semrock, Buffalo NY, USA). Emission was passed through a 536/40 nm filter (Semrock) and recorded with a photomultiplier (H9656-20; Hamamatsu Photonics, Hamamatsu, Japan). Signals were amplified with a lock-in amplifier (MFLI, Signal Zürich Instruments, Switzerland) and recorded with a data acquisition pad (PCI-6221; National Instruments, Germany) and BioKine software v. 4.49 (Bio-Logic).

### Electrophysiology

We recorded CatSper currents from human sperm in the whole-cell configuration, as described before (Strunker et al., 2011). Seals between pipette and sperm were formed either at the cytoplasmic droplet or in the neck region in solution (HS) containing (in mM): 135 NaCl, 5 KCl, 1 MgSO_4_, 2 CaCl_2_, 5 glucose, 1 Na-pyruvate, 10 lactic acid, 20 HEPES, adjusted to pH 7.4 with NaOH. Monovalent CatSper currents were recorded in a sodium-based divalent-free solution (NaDVF) containing (in mM): 140 NaCl, 40 HEPES, and 1 EGTA, adjusted to pH 7.4 with NaOH. The osmolarity of HS and NaDVF solution was approximately 320 mOsm. The pipette (10–15 MΩ) solution contained (in mM): 130 Cs-aspartate, 50 HEPES, 5 EGTA, 5 CsCl, adjusted to pH 7.3 with CsOH. The pipette solution was approximately 325 mOsm. To examine the effect of intracellular cAMP on CatSper, cAMP was dissolved in the pipette solution. The action of extracellular application of H89 (50 mM stock in DMSO), KT 5720 (20 mM stock in DMSO), PKI 14-22 (800 μM stock in water) was examined by diluting the stocks with NaDVF; 8-CPT-cAMP was directly dissolved at 5 mM in NaDVF. All experiments were performed at room temperature (21-25°C). Data were not corrected for liquid junction potentials.

### Measurement of intracellular cAMP content

To examine the action of adenosine and IBMX, 308 µl of sperm in HTF at a density of 2 × 10^7^ cells/ml (6 × 10^6^ cells in total) were mixed with 32 µl of HTF containing adenosine or IBMX to reach a final concentration of (in mM): 0.1 adenosine and 0.5 IBMX. To examine the action of bicarbonate, 154 µl of sperm in bicarbonate-free HTF at a density of 4 × 10^7^ cells/ml (6 × 10^6^ cells in total) were mixed with 154 µl HTF containing 46 mM NaHCO_3_ and 32 µl of HTF containing 25 mM NaHCO_3_. After mixing with the respective stimulus, the samples were incubated for 30 min at 37°C, followed by the addition of 18 µl of 5 M HCl (0.25 M final concentration) to quench the biochemical reactions. After incubation for 30 min at room temperature, cell debris was sedimented by centrifugation at 3,000 g for 5 min at room temperature. The cAMP concentration in the supernatant was determined by a competitive enzyme immunoassay according to the product manual (Catalog #: ADI-900-066, Enzo Life Sciences), including acetylation of cAMP. Calibration curves were obtained by serial dilutions of cAMP standards (acetylated format).

### Data analysis

All data are given as mean ± SD.

## Acknowledgments

We thank Sabine Forsthoff and Jolanta Körber-Naprodzka for technical support and U. Benjamin Kaupp and Dagmar Wachten for critical reading of the manuscript.

## Author contributions

T.S., C.B., X.H.Z., and T.W. conceived the project. All authors designed research, performed experiments, acquired, analyzed, and/or interpreted data. T.S., C.B., and T.W. wrote the manuscript. All authors revised the manuscript critically for important intellectual content, and approved the manuscript.

## Data and materials availability

All data are present in the main text, figures, and the supplementary material.

---

**Supplementary Figure 1.**
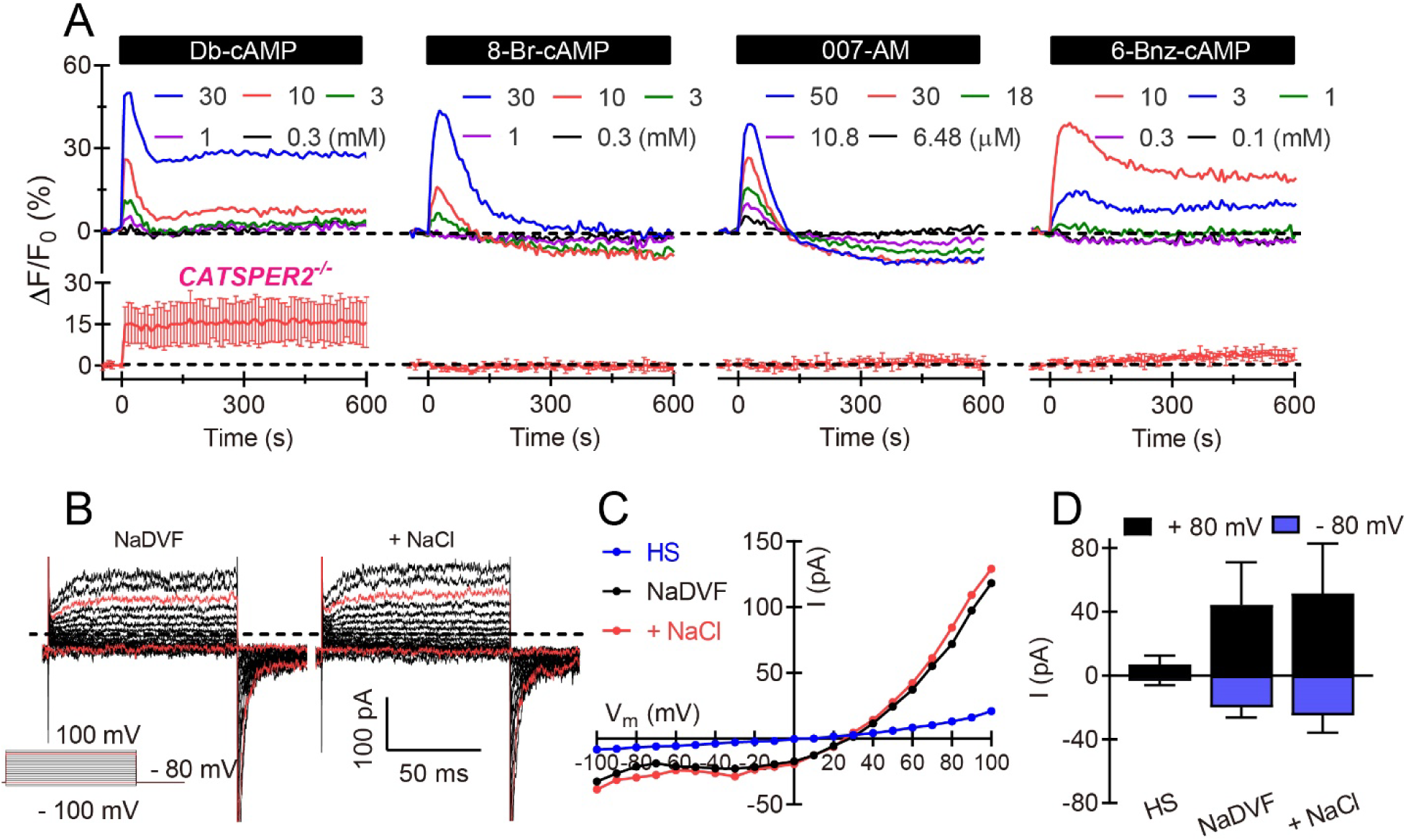
cAMP analogs evoke Ca^2+^ signals in human sperm. **(A)** Representative Ca^2+^ signals evoked by Db-cAMP, 8-Br-cAMP, 8-pCPT-2-O-Me-cAMP (e.g. 007-AM), and 6-Bnz-cAMP in sperm from healthy donors (upper panel), and averaged Ca^2+^ signal (lower panels, n = 3) in sperm lacking functional CatSper channels (*CATSPER2^-/-^*). **(B)** Representative whole-cell currents at pH_i_ 7.3 in divalent-free Na^+^-based bath solution (NaDVF) and in NaDVF containing additional 5 mM NaCl. Dotted black line: zero current level. Red traces: currents at +80 mV and −80 mV. **(C)** Steady-state current-voltage relationship from (B) and in extracellular solution containing Ca^2+^ and Mg^2+^ (HS) indicated as a control. **(D)** Mean current amplitudes at +80 mV and −80 mV in HS, in NaDVF, and in NaDVF containing additional 5 mM NaCl (n = 3).

**Supplementary Figure 2.**
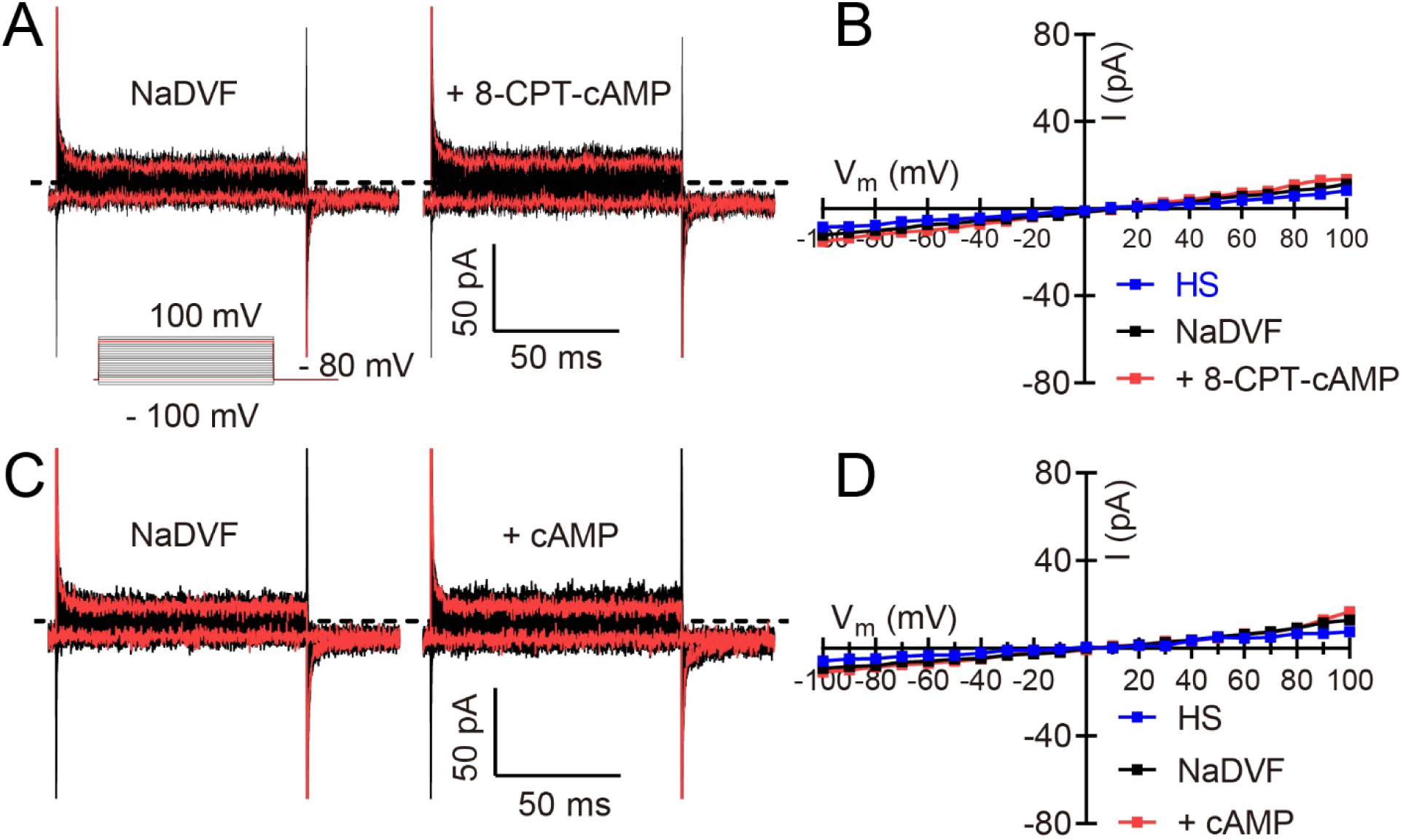
Action of 8-CPT-cAMP and cAMP in CatSper-deficient human sperm. **(A, C)** Representative whole-cell currents at pH_i_ 7.3 recorded from sperm lacking functional CatSper channels *(CATSPER2^-/-^*) in divalent-free Na^+^-based bath solution (NaDVF) and in NaDVF containing 5 mM 8-CPT-cAMP (A) or 10 mM cAMP (C). Dotted black line: zero current level. Red traces: currents at +80 mV and −80 mV. **(B, D)** Steady-state current-voltage relationship from (A) and (C), respectively, with currents in extracellular solution containing Ca^2+^ and Mg^2+^ (HS) indicated as a control.

**Supplementary Figure 3.**
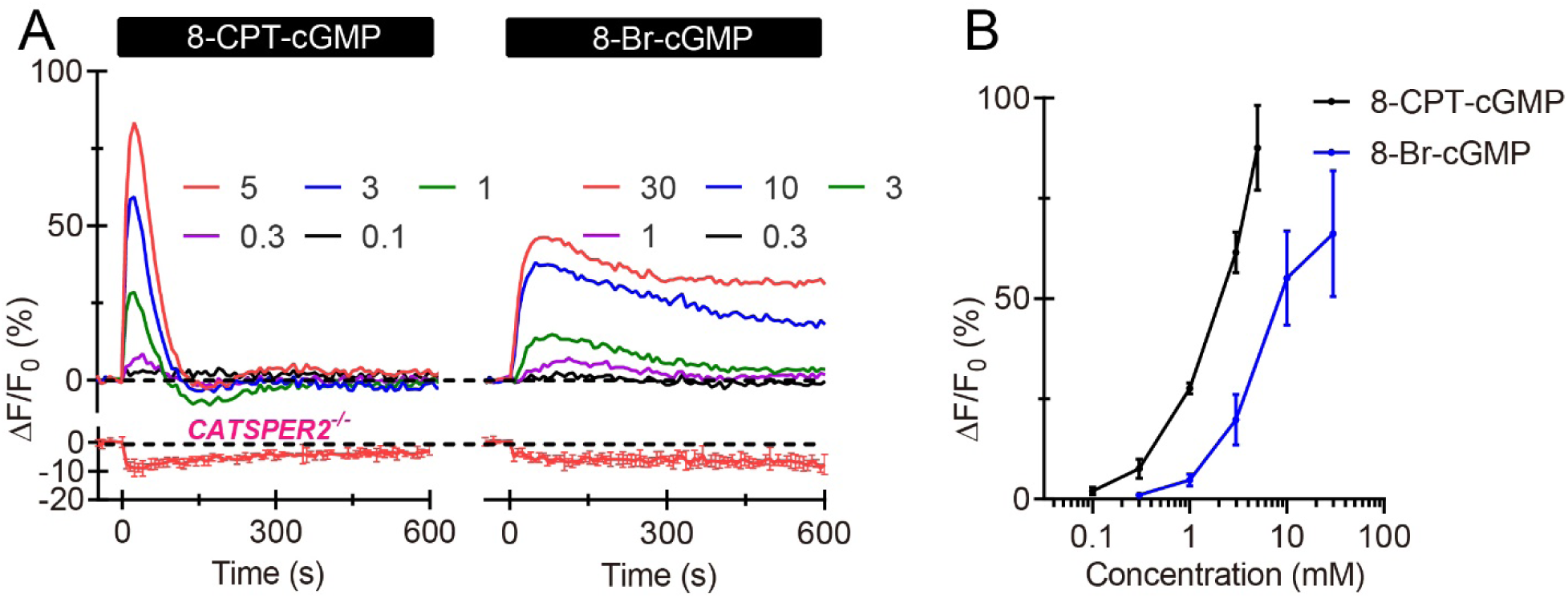
cGMP analogs activate CatSper in human sperm. **(A)** Representative Ca^2+^ signals evoked by 8-CPT-cGMP and 8-Br-cGMP in sperm from healthy donors and averaged Ca^2+^ signal (lower panels, n = 3) in sperm lacking functional CatSper channels (*CATSPER2^-/-^*). **(B)** Mean amplitudes of Ca^2+^ signals evoked by 8-CPT-cGMP and 8-Br-cGMP (n = 3) in sperm from healthy donors.

**Supplementary Figure 4.**
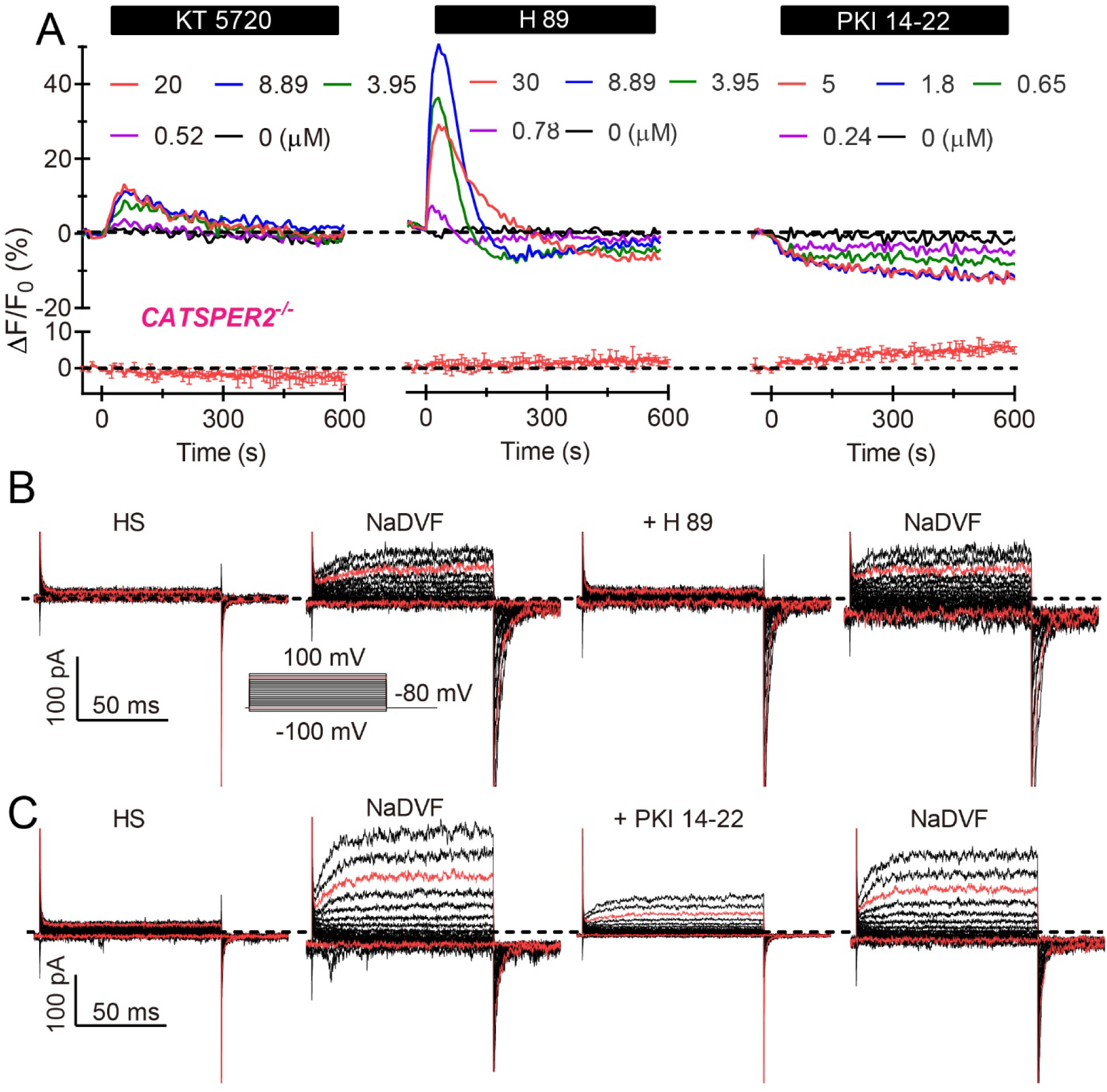
The actions of PKA inhibitors in human sperm. **(A)** Representative Ca^2+^ signals evoked by KT 5720, H89, and PKI 14-22 in sperm from healthy donors and averaged Ca^2+^ signal (lower panels, n = 3) in sperm lacking functional CatSper channels (*CATSPER2^-/-^*). **(B and C)** Representative whole-cell currents at pH_i_ 7.3 in extracellular solution containing Ca^2+^ and Mg^2+^ (HS), in divalent-free Na^+^-based bath solution (NaDVF), and in NaDVF containing 30 μM H89 (B) or 5 µM PKI 14-22 (C). Dotted black line: zero current level. Red traces: currents at +80 mV and −80 mV.

**Supplementary Figure 5.**
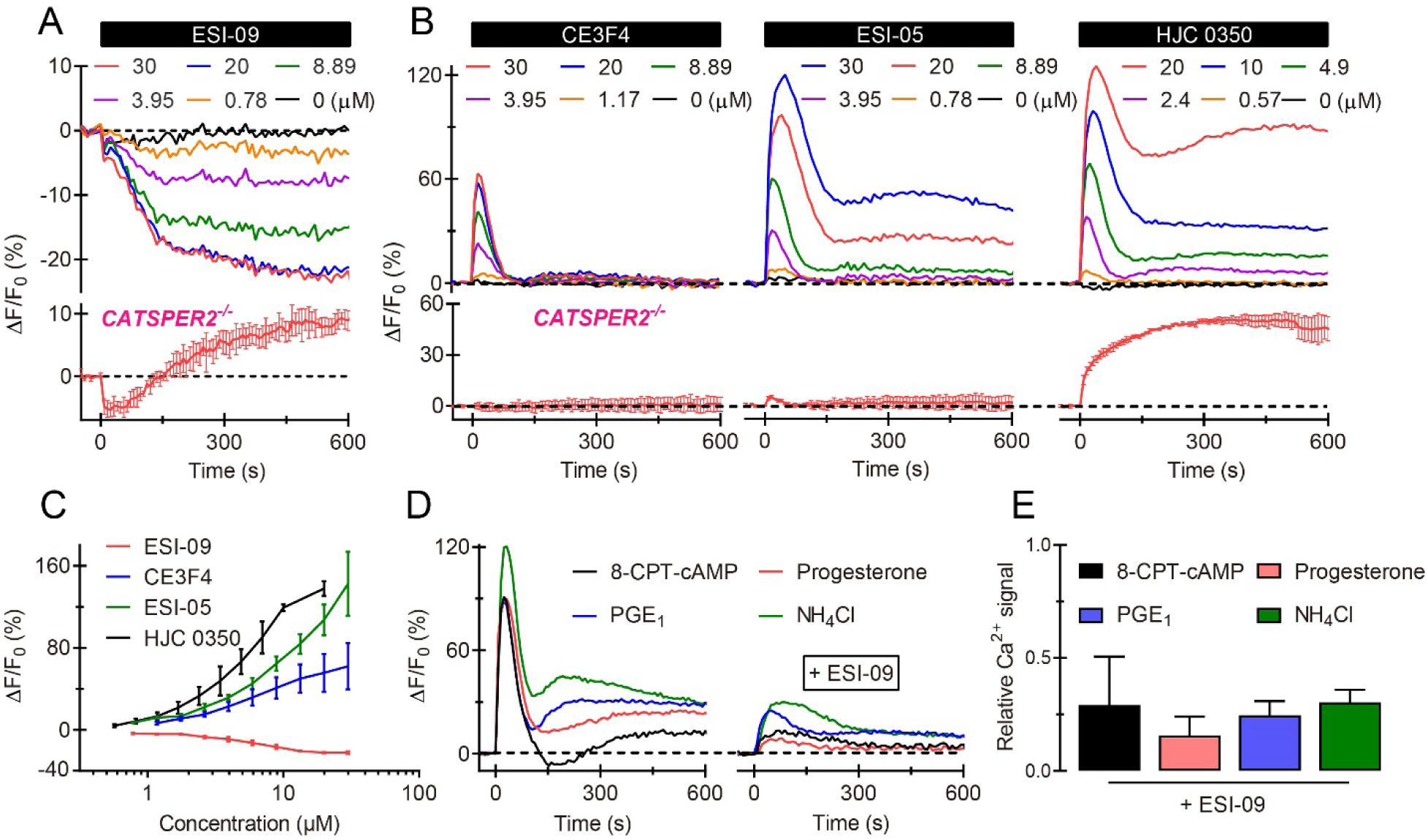
The action of EPAC inhibitors in human sperm. **(A, B)** Representative Ca^2+^ signals evoked ESI 09, CE3F4, ESI 05, and HJC 0350 in sperm from healthy donors and averaged Ca^2+^ signal (lower panels, n = 3) in sperm lacking functional CatSper channels (*CATSPER2^-/-^*). **(C)** Mean amplitudes of Ca^2+^ signals evoked by EPAC inhibitors (n ≥ 4). **(D)** Ca^2+^ signals evoked by 8-CPT-cAMP (5 mM), progesterone (10 μM), PGE_1_ (10 μM), and NH_4_Cl (10 mM), in the absence and presence of ESI 09 (30 μM). **(E)** Mean amplitude of Ca^2+^ signals in the presence of ESI 09 (30 μM) relative the amplitude evoked in the absence of any drug (n ≥ 3).

